# Identification of antibiotic resistance genes in fecal microbiota selected donors during the establishment of a biobank in the south of Brazil

**DOI:** 10.64898/2026.05.07.723634

**Authors:** Lucas de Figueiredo Soveral, Liandra Raphaella de Lima Holanda, Izadora Borgmann Frizzo, Leonardo Gonçalves Gomes, Isabella Bittencourt de Souza, Gabriella de Souza, Patrícia Almeida Vanny, Oscar Bruna-Romero, Jussara Kasuko Palmeiro, Mara Cristina Scheffer, Thais Cristine Marques Sincero, Carlos Rodrigo Zárate-Bladés

## Abstract

Fecal microbiota transplantation (FMT) is an effective therapy for recurrent Clostridioides difficile infection and is increasingly explored for other dysbiosis-related disorders. However, its implementation as a regulated therapeutic strategy still requires robust donor screening, biosafety frameworks, and standardized processing workflows. Here, we describe the establishment of the first fecal microbiota biobank in the south of Brazil and evaluate the incorporation of metagenomic sequencing as a complementary layer of donor safety assessment. A structured donor selection pipeline based on international guidelines was implemented, integrating clinical screening, biochemical and serological testing, and microbiological analyses. Of 100 screened candidates, only four donors met all eligibility criteria and were included in the biobank, highlighting the stringency of the selection process. Shotgun metagenomic sequencing revealed a diverse resistome across all donors, including a shared core set of resistance-related genes alongside marked interindividual variability. Dominant antibiotic resistance genes included tetracycline-associated determinants, as well as *ermF, CfxA*-type β-lactamases, and aminoglycoside-modifying enzymes, each linked to specific gut taxa. Notably, the relatively high abundance of *tetW* and *ermF* in *Bacteroides fragilis* suggests that this dominant commensal species may act as a reservoir for tetracycline and multidrug resistance determinants within the intestinal microbiota. Rather than serving as exclusion criteria, such determinants highlight the importance of integrating functional genomic profiling into donor characterization. Overall, this study provides a framework for microbiota biobank implementation and supports the use of metagenomics as a complementary strategy to improve biosafety and functional assessment in FMT.

## 1. Introduction

Fecal microbiota transplantation (FMT) has consolidated itself as one of the most effective therapeutic interventions for recurrent *Clostridioides difficile* infection, with success rates exceeding 90% (1,2). Beyond this indication, FMT has progressively gained attention as a potential therapeutic strategy for a wide spectrum of disorders linked to intestinal dysbiosis, including inflammatory, metabolic, and immune-mediated diseases (3–5).

Despite its clinical relevance, FMT poses singular challenges from a technical and safety perspective. Unlike conventional pharmaceuticals or biologics composed of defined molecular entities, FMT involves the transfer of a complex and dynamic biological ecosystem, whose composition and functional attributes remain only partially characterized (6–8). This intrinsic complexity complicates standardization, traceability and quality control, fundamental requirements for the development of safe and scalable therapeutic products (9,10).

In this context, fecal microbiota biorepositories and stool banks have emerged as essential infrastructures for the consolidation of FMT as a regulated therapy. These platforms aim to centralize donor recruitment, ensure standardized screening, enable controlled processing and storage, and support the traceability of material used for clinical application (11,12). However, the implementation of such systems remains heterogeneous across countries, reflecting differences in regulatory environments, technological capacity, and clinical practice (9,13,14).

Current donor screening protocols are primarily designed to exclude known pathogens and high-risk clinical conditions (15). While indispensable, these strategies are limited to predefined targets and do not fully encompass the functional and genetic complexity of donor microbiota (16,17). As FMT expands toward new clinical indications and increasingly vulnerable patient populations, the need for more comprehensive and function-oriented approaches to biosafety becomes increasingly evident (18,19).

Within this, the present study describes the establishment of the first fecal microbiota Biobank in Santa Catarina and explore the use of metagenomic sequencing as a complementary tool for safety assessment, not as a definitive safety criterion, but as a way for expanding quality control strategies in microbiota-based therapies.

## 2. Materials and Methods

### 2.1. Study design

This study was designed as a descriptive and methodological investigation aimed at establishing the first fecal microbiota biobank in the state of Santa Catarina, Brazil, intended to support the future clinical use of fecal microbiota transplantation (FMT) in selected diseases. The study was structured in three main sequential phases: (i) development of donor screening tools based on international guidelines; (ii) implementation of a clinical and laboratory-based donor selection pipeline; and (iii) exploratory incorporation of metagenomic analysis of donor’s fecal microbiota.

### 2.2. Development of donor screening questionnaire and testing pipeline

Initially, a comprehensive literature review was performed to identify and compare current recommendations for FMT donor selection and biosafety. This review focused on three core components: clinical exclusion criteria, blood and serological testing, and microbiological screening of fecal samples. The screening tests developed by major national and international working groups were analyzed and compared, including those from the IAG HC-UFMG (14), the Netherlands Working Group (20), the International Consensus Conference on FMT (21), the European FMT Working Group (9), and the Australian Consensus (13). Based on this comparative analysis and additional literature, a customized screening questionnaire and testing pipeline were developed and implemented in the present work (Supplementary Tables 1 and 2).

### 2.3. Donor recruitment and clinical screening

Donor recruitment was conducted between July and September 2023 through public calls and institutional communication channels. Eligible volunteers were invited to complete a structured clinical screening questionnaire based on international FMT guidelines. Candidates presenting one or more exclusion criteria were not allowed to proceed to subsequent screening steps. Exclusion criteria included, but were not limited to, recent antibiotic use, chronic gastrointestinal diseases, metabolic or autoimmune disorders, recent infectious diseases, high-risk behaviors, and abnormal bowel habits.

### 2.4. Laboratory tests

Candidates who met the clinical eligibility criteria were submitted to a comprehensive laboratory screening protocol, designed to assess general health status and exclude infectious agents potentially transmissible through fecal microbiota transplantation. Blood-based analyses included a complete blood count, serum electrolytes (sodium, potassium, calcium, magnesium), renal and hepatic function markers (urea, creatinine, AST/TGO, ALT/TGP, GGT, bilirubin), metabolic and nutritional parameters (glucose, vitamin B12, folic acid, vitamin D), and inflammatory markers (C-reactive protein). Serological screening comprised tests for HIV types I and II, hepatitis A (IgM/IgG), hepatitis B (HBsAg and anti-HBc IgG/IgM), hepatitis C (anti-HCV), human T-lymphotropic virus (HTLV I/II), cytomegalovirus (IgG/IgM), and Epstein–Barr virus (IgG/IgM).

Fecal samples were analyzed for enteric pathogens using conventional microbiological methods and molecular assays. These included coproculture and PCR-based detection of *Clostridioides difficile* toxin A/B, rotavirus, adenovirus, SARS-CoV-2, herpes simplex virus I and II (IgG and IgM), and a multiplex PCR panel (FilmArray®) targeting bacterial, viral, and parasitic pathogens, such as *Campylobacter* spp., *Salmonella* spp., *Shigella*/EIEC, *Vibrio cholerae, Yersinia enterocolitica, Clostridioides difficile, Giardia lamblia, Entamoeba histolytica, Cryptosporidium, Cyclospora cayetanensis*, norovirus GI/GII, astrovirus, sapovirus, and adenovirus F40/41. Only candidates presenting normal results across all clinical, serological, biochemical, and microbiological tests were considered eligible donors for inclusion in the biobank.

### 2.5. Processing and storage of fecal material

Fecal material from eligible donors was processed under controlled laboratory conditions following standardized operating procedures. Briefly, samples were collected in clean conditions using and sterile recipient. Next, samples were homogenized, filtered to remove particulate debris, aliquoted, and cryopreserved at ultra-low temperatures. All samples were labeled and documented to ensure traceability, and storage conditions were continuously monitored.

### 2.6. Antibiotic resistance genes identification by shotgun metagenomics

Total genomic DNA from fecal samples obtained from eligible donors was extracted using the Quick-DNA™ HMW MagBead Kit (Zymo Research, Irvine, CA, USA), following the manufacturer’s protocol optimized for recovery of high-molecular-weight DNA from complex biological matrices. DNA quantity was measured using the Qubit fluorometer (Thermo Fisher Scientific, Waltham, MA, USA), and purity was evaluated by spectrophotometry using the NanoVue system (GE Healthcare, Chicago, IL, USA). Metagenomic sequencing libraries were prepared using the Rapid Barcoding Kit 24 v14 (SQK-RBK114.24; Oxford Nanopore Technologies, Oxford, UK), according to the manufacturer’s instructions. Barcoded libraries were pooled and sequenced for 48 hours on a MinION platform (Oxford Nanopore Technologies) using R10.4.1 flow cell chemistry. Raw sequencing reads were basecalled, demultiplexed, and quality-filtered using Oxford Nanopore standard workflows and Guppy v4.3.0. Reads passing quality control were used for downstream resistome analysis.

### 2.7. Data Analysis

Antibiotic resistance determinants were identified by alignment against the Comprehensive Antibiotic Resistance Database (CARD). Due to the exploratory nature of the study, no statistical analyses were performed. Gene presence/absence, relative abundance, and cumulative abundance profiles were subsequently used for comparative analyses across samples using R™ (Version 4.3.2).

### 2.8. Ethical Considerations

The study was approved by the institutional review board under process number CAAE: 42603020.0.0000.0121. All participants provided written informed consent prior to sample collection and laboratory analyses.

## 3. Results

### 3.1. Donor recruitment and clinical eligibility

A total of 100 candidates were included in the clinical screening questionnaire during the recruitment period. Of these, 6 candidates met the clinical eligibility criteria and were referred to laboratory screening. After biochemical, serological, and microbiological analyses, another two additional candidates were excluded, and only four participants were ultimately considered eligible for fecal microbiota donation and inclusion in the biobank. The donor selection workflow and the main exclusion criteria identified are illustrated in Figure 1.

**Figure 1.**
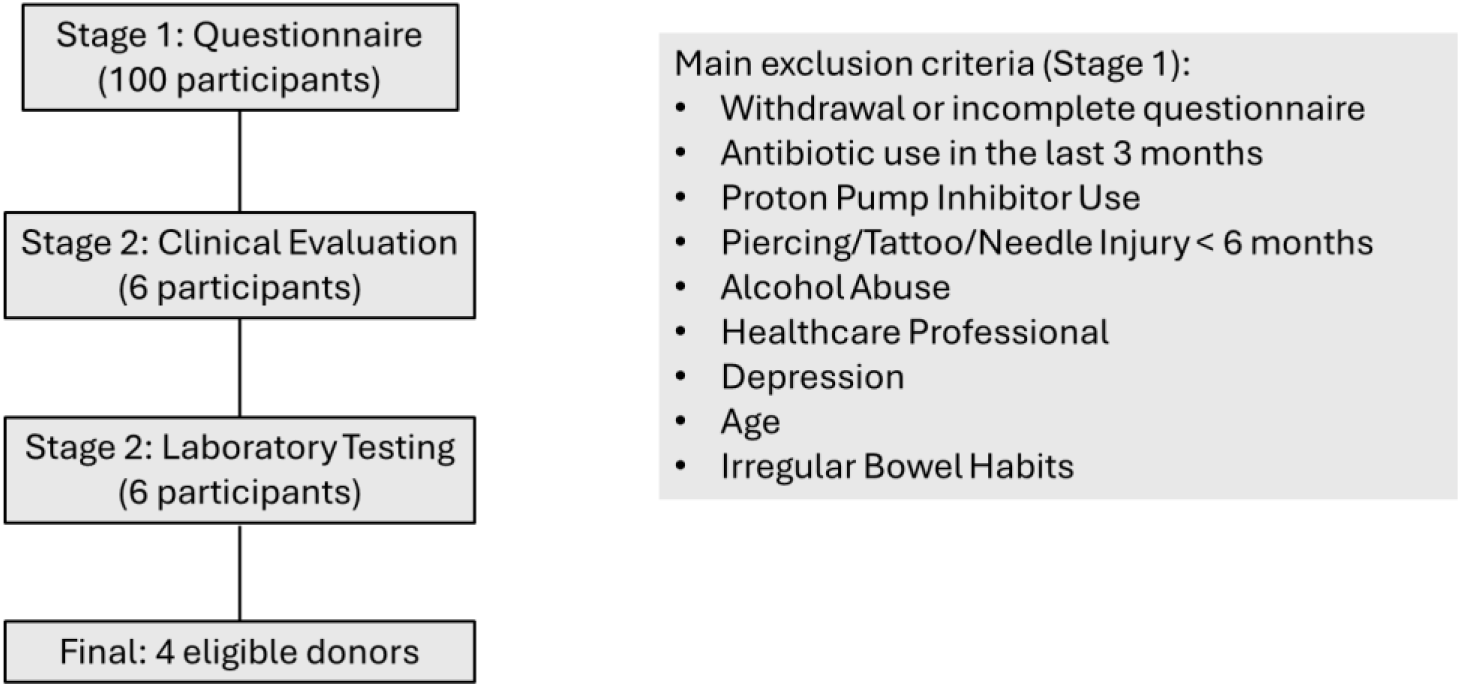
Flowchart illustrating the multistep donor selection process.

### 3.2. Resistome profile characterization

Sequencing generated 7.09 Gb of data across all samples (5.44 million reads; N50 = 2.55 kb), with read accuracy ranging from 79.6% to 84.7%, indicating overall good sequencing performance. Shotgun metagenomic analysis of fecal samples from eligible donors identified 180 different resistance-related gene entries in CARD dataset. Detection frequency varied between samples, with 137 genes identified in Sample 1, 87 in Sample 2, 103 in Sample 3, and 134 in Sample 4. Of these, only 58 were consistently detected in all samples, whereas 122 showed partial occurrence in different samples (Figure 2 A). To improve biological interpretability, detected entries were grouped into two major categories: ARGs, comprising canonical resistance genes whose presence is directly associated with antimicrobial resistance phenotypes (n= 100), and variant-dependent loci, corresponding to conserved chromosomal targets in which resistance is typically mediated by specific sequence variants rather than gene presence alone (e.g., rRNA genes, gyrA/parC/parE, and rpoB-associated loci) (n=80). Among consensus genes, only eleven were classified as canonical ARGs (Core resistome), including APH(3’)-IIIa, CfxA2, CfxA3, ErmF, mefA, AlaS and five tetracycline resistance determinants (tet(32), tet(40), tet(O), tet(Q), and tet(W)).

**Figure 2.**
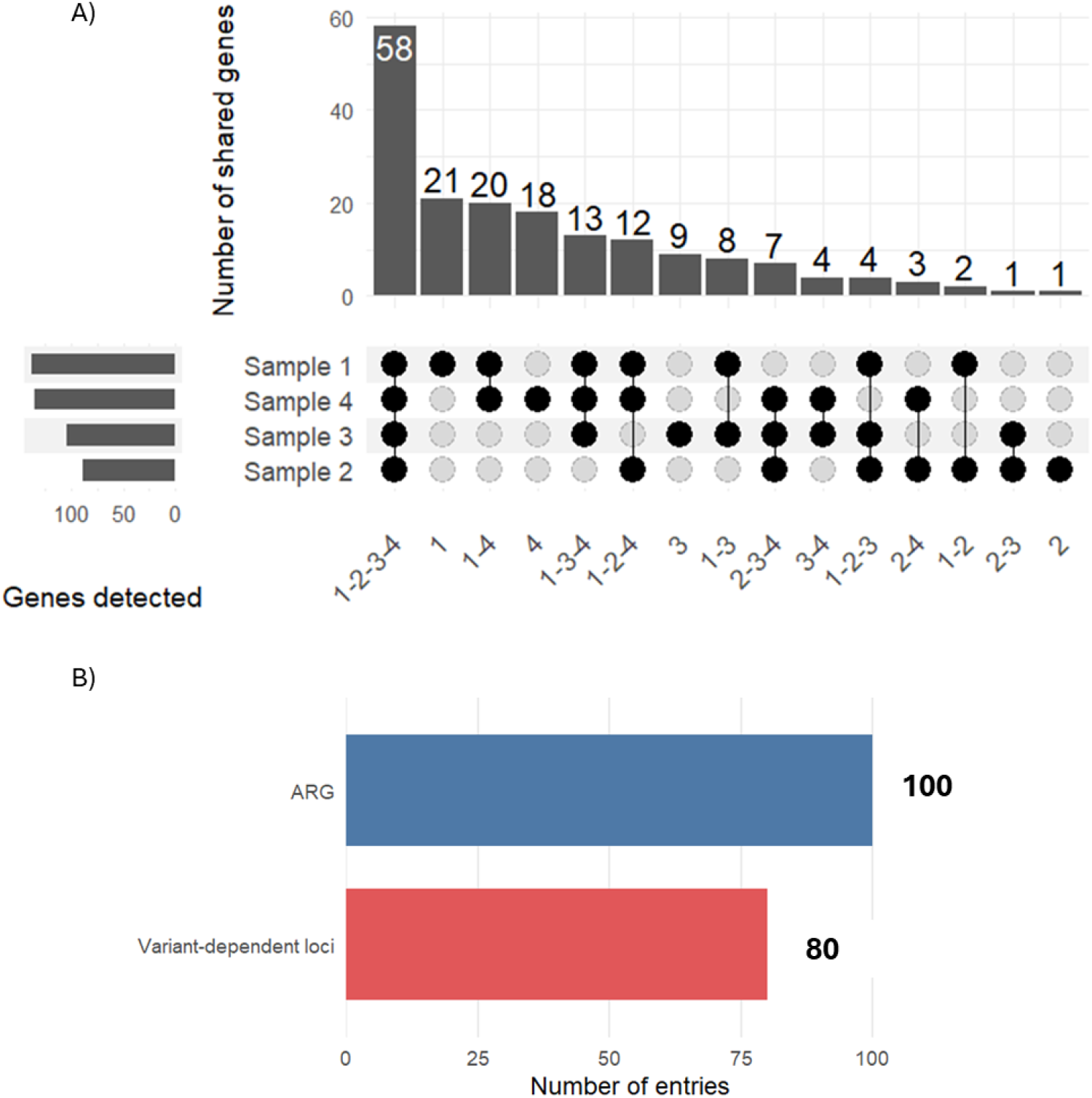
UpSet plot showing the intersection (number) of resistance-related genes detected across the four samples. The bar plot (top) indicates the number of genes shared among specific combinations of samples, while the dot matrix (bottom) represents the corresponding sample intersections. A total of 58 genes were detected in all four samples, while additional genes were shared among subsets of samples or were sample-specific **(A).** Horizontal bars (left) indicate the total number of genes detected in each individual sample. Classification of resistance-related entries according to functional category **(B)**.

### 3.3. Taxonomic distribution and abundance of dominant antibiotic resistance genes

The taxonomic distribution of the most abundant ARGs across samples is shown in Figure 3A. A consistent set of dominant resistance genes was observed across all samples, with tet(Q), tet(W), tet(O), ErmF, and CfxA2 representing the most prominent determinants. These genes were strongly associated with specific bacterial taxa, indicating clear links between resistome composition and microbiota structure. In particular, tet(Q) was predominantly associated with *Bacteroides fragilis*, while tet(W) was mainly linked to *Bifidobacterium longum*. The gene tet(O) was primarily associated with *Campylobacter jejuni*, whereas ErmF was also largely attributed to *Bacteroides fragilis*. In contrast, CfxA2 was associated with *Prevotella intermedia*, highlighting additional taxonomic contributors to β-lactam resistance.

**Figure 3.**
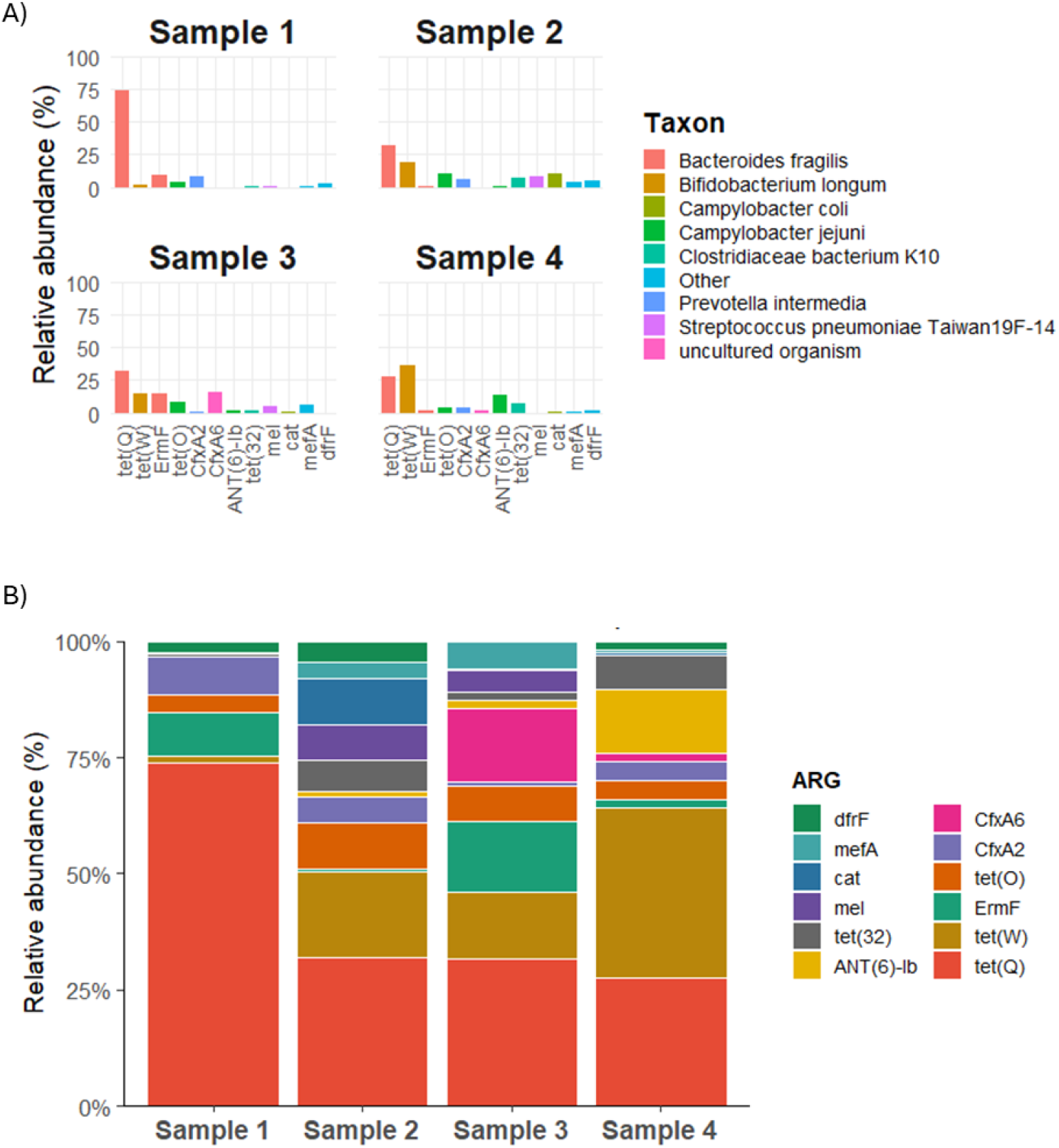
**(A)** Individual bar plots for each sample showing the distribution (%) of the 12 most abundant resistance genes, categorized by their associated bacterial taxa. The color legend indicates taxonomic group carrying the detected ARGs. **(B)** Stacked bar plot illustrating the distribution of 12 most abundant ARGs across all samples. Each color represents a specific ARG.

Although these genes were detected across all samples, their proportional abundance varied markedly (Figure 3 B). Sample 1 was strongly dominated by tet(Q), which accounted for 34.3% of ARG abundance, followed by ErmF (4.4%) and APH(3’)-IIIa (1.5%). In Sample 2, a more distributed profile was observed, with tet(Q) (9.1%), tet(O) (2.9%), tet(32) (1.9%), and tet(40) (1.7%) contributing to the resistome, along with macrolide resistance genes such as mel (2.2%). Sample 3 showed a more balanced composition, with tet(Q) (15.2%), ErmF (7.3%), tet(W) (7.0%), and CfxA6 (7.6%) representing major contributors. In Sample 4, the dominance of tet(Q) was reduced (4.6%), with increased contributions from tet(W) (6.1%) and ANT(6)-Ib (2.3%), indicating a shift toward a more heterogeneous resistome structure.

### 3.4. Distribution of resistance classes and mechanisms

The resistome was predominantly composed of genes associated with aminoglycoside resistance (32.9%) and tetracycline resistance (28.9%), which together accounted for more than half of the total ARG abundance. Genes related to multidrug resistance also represented a substantial fraction (19.3%), indicating the presence of broad-spectrum resistance determinants. Other classes were detected at lower frequencies, including cephalosporin (5.1%), peptide (3.4%), and elfamycin (2.3%), while macrolide (1.2%) and streptogramin (1.1%) resistance genes contributed minimally to the overall resistome.

The functional mechanisms underlying resistance are summarized in Figure 4B. The resistome was strongly dominated by target alteration mechanisms (58.2%), followed by target protection (26.8%), indicating that resistance in these samples is primarily mediated through modifications of antibiotic targets rather than drug degradation or efflux. Enzymatic inactivation mechanisms accounted for 8.4%, while efflux systems contributed 2.9% of the total ARG abundance. Additional mechanisms, including target replacement (1.2%) and unclassified categories (2.4% unknown; 0.1% other), were detected at low frequencies.

**Figure 4.**
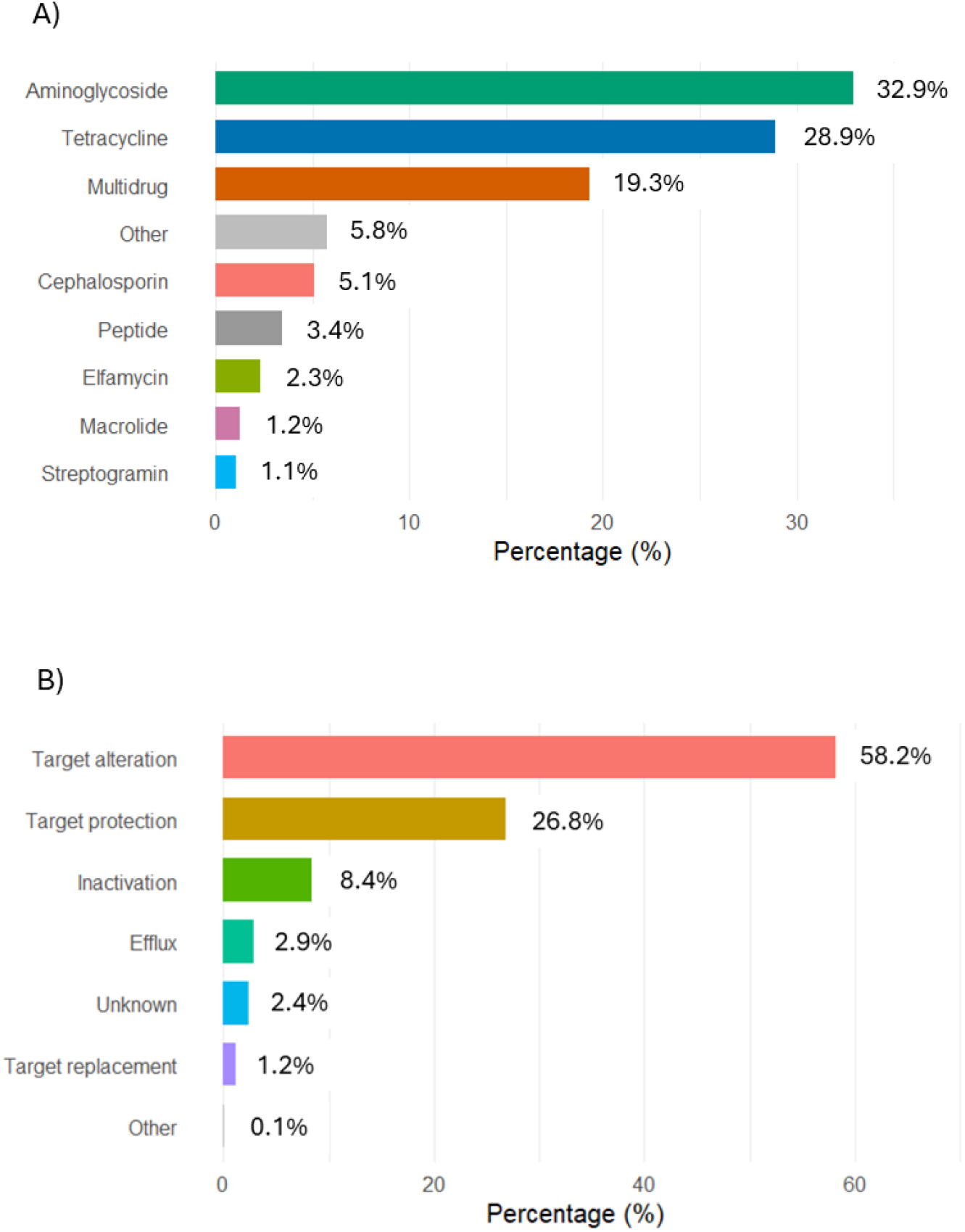
Distribution of resistance genes according to the class of antibiotics **(A)**. Relative abundance of antimicrobial resistance mechanisms identified across the samples **(B)**.

## 4. Discussion

A central challenge in FMT implementation worldwide remains the identification of suitable donors. Our findings confirm that donor selection is a highly restrictive process, with only a minority of screened candidates ultimately qualifying for donation. Similar donor approval rates have been reported in international biobanking initiatives, reflecting the stringency required to minimize transmissible risks while preserving microbiota diversity and therapeutic potential (9,14). This reinforces the importance of structured, transparent, and reproducible selection workflows as a foundational element of any FMT program. By integrating clinical questionnaires, biochemical and serological testing, and microbiological screening, our workflow aligns with major international guidelines while being operationally feasible within public and private healthcare infrastructures in Brazil (21). This aspect is particularly relevant in low- and middle-income countries, where the translation of advanced microbiota-based therapies often faces infrastructural and regulatory barriers.

An important contribution of this study is the incorporation of metagenomic sequencing and identification of ARGs as a complementary layer in donor safety assessment. As expected, tetracycline resistance determinants such as *tet(Q)* and *tet(W)* were highly prevalent and primarily associated with *Bacteroides fragilis* and *Bifidobacterium longum*, respectively, reinforcing the well-established role of these taxa as major reservoirs of tetracycline resistance in the human gut (22,23). Notably, the relatively high abundance of *ermF*, also linked to *Bacteroides fragilis*, a dominant commensal bacterium, suggests that this species may act as a reservoir for multidrug resistance determinants. In addition, *CfxA2/6* and *ANT(6)-Ib*, associated with *Prevotella* and *Campylobacter*, respectively, further highlight the contribution of gut commensals to β-lactam and aminoglycoside resistance pools. These genes were not uniformly distributed across samples, but instead showed distinct abundance patterns, with *tet(Q)* dominating in some donors and a more balanced ARG composition observed in others. At the functional level, aminoglycoside, tetracycline, multidrug, and cephalosporin resistance classes accounted for most of the ARG abundance. Consistently, resistance mechanisms were predominantly driven by target alteration and target protection, along with a smaller contribution from enzymatic inactivation.

In a clinical perspective, seminal evidence provided by DeFilipp et al. demonstrated that under specific high-risk conditions FMT can mediate the transmission of multidrug-resistant organisms, including extended-spectrum β-lactamase (ESBL)-producing *Escherichia coli*, with severe clinical consequences in immunocompromised recipients (24). In this study, genomic sequencing traced bacteremia-causing strains back to donor material, highlighting that conventional donor screening may fail to detect clinically relevant resistance determinants and that host factors such as impaired gut barrier integrity, neutropenia, and concurrent antibiotic exposure critically modulate risk. In our results, the relatively high abundance of *ermF*, a multidrug resistance gene associated with the MLS_B_ phenotype (conferring resistance to macrolides, lincosamides, and streptogramin B) in *Bacteroides fragilis* raises the possibility that such genes may serve as reservoirs for horizontal gene transfer under selective pressure, particularly in vulnerable hosts (23).

In contrast Woodworth *et al* indicates that fecal microbiota transplantation may also actively reduce the burden of resistance genes and multidrug-resistant organisms in the gut (25).

In this work, FMT led to sustained decolonization of multi-drug resistant organisms (MDRO) in the majority of treated patients and significantly prolonged the time to recurrent MDRO infection when compared to untreated controls. This work shows that susceptible Enterobacterales strains replace baseline Extended-Spectrum β-Lactamase-producing strains in recipients, suggesting that FMT can reshape the intestinal ecosystem in a way that favors the expansion of less resistant endogenous populations.

Taken together, the detection of antimicrobial resistance–associated genes in clinically eligible donors should not be interpreted as a marker of donor inadequacy or direct clinical risk. However, considering the potential for horizontal transfer of resistance determinants, these genes should not be overlooked in safety assessments, particularly when FMT is intended for highly vulnerable recipients, such as immunocompromised or critically ill patients.

Moreover, these observations further support the exploration of alternative FMT formulations, such as sterile fecal filtrates or defined microbial consortia, which do not rely on the transfer of viable bacterial communities and may offer enhanced biosafety profiles (26,27). Early clinical evidence suggests that some of these approaches can achieve therapeutic efficacy comparable to conventional FMT, although robust validation in larger and diverse clinical settings is still required (26,28). Overall, strategies that simultaneously enhance biosafety, improve standardization, and facilitates donor selection should be actively considered as the field moves toward more refined and scalable microbiota-based therapies.

## 5. Limitations and Perspectives

Some limitations of this study should be acknowledged. First, the number of participants who completed the full donor selection pipeline was relatively small, as well as the number of samples subjected to metagenomic sequencing, which restricts a more robust comparative analysis. Second, the resistome characterization was exploratory and not designed to establish clinical risk or to redefine biosafety criteria for fecal microbiota transplantation.

Finally, the use of a single sequencing platform and analytical pipeline may introduce methodological biases inherent to any metagenomic approach, which should be considered when interpreting the results. Despite these limitations, the present work provides evidence for integrating metagenomic tools into microbiota biobank development and supports future studies aimed at expanding functional characterization layers, increasing donor cohorts, and correlating microbiota profiles with clinical efficacy and safety in microbiota-based therapies.

## Author Contributions

L.F.S. contributed to the development of the donor screening workflow, performed metagenomic and resistome data analyses, prepared the figures, interpreted the data, and wrote the manuscript. L.R.L.H. contributed to donor screening procedures, sample collection for laboratory testing, and sequencing procedures. I.B.F. performed sequencing procedures and contributed to the analysis of sequencing data. L.G.G. contributed to donor screening procedures, sample collection, and sequencing procedures. I.B.S. contributed to sample collection and sequencing procedures. G.S. contributed to data analysis. P.A.V. provided technical support. O.B.R., J.K.P., M.C.S., and T.C.M.S. contributed with methodological and scientific support. C.R.Z.B. conceived and supervised the study, contributed to data interpretation, and critically revised the manuscript. All authors reviewed and approved the final manuscript.

## Ethics Statement

Approved under process number CAAE: 42603020.0.0000.0121.

