## Supplementary Table 1 for "Identification of antibiotic resistance genes in fecal microbiota selected donors during the establishment of a biobank in the south of Brazil"

Supplementary Table 1. Comparison of exclusion criteria for fecal microbiota transplantation donor screening across international guidelines

| Exclusion Criteria | UFMG (2020) | Amsterdam (2013) | International (2019) | European (2016) | Australian (2020) |
|---|---|---|---|---|---|
| Known uncontrolled systemic infection at the time of donation | X | | X | X | |
| Use of illicit drugs | X | X | X | X | X |
| High-risk sexual behavior (anonymous partners; contact with sex workers, drug users, or individuals with HIV, viral hepatitis, or syphilis; sex work; history of sexually transmitted infections) | X | X | X | X | X |
| Previous receipt of organ/tissue transplant | X | X | X | X | |
| Previous receipt of blood products | <6 months | X | | <12 months | |
| Recent needlestick injury | <6 months | X | | <6 months | <3 months |
| Recent tattoo, piercing, earrings, acupuncture | <6 months | X | <6 months | <6 months | <3 months |
| Recent parasitic infection or gastrointestinal infection (e.g., rotavirus, *Giardia lamblia*) | | | ≤2 months | X | |
| Recent travel (<6 months) to tropical countries or regions with high risk of infectious diseases or traveler's diarrhea | | X | X | X | <3 months |
| Recent vaccination with live attenuated virus (if potential transmission risk exists) | <2 months | | ≤2 months | <6 months | |
| Healthcare workers (risk of multidrug-resistant organism transmission) | X | X | | X | |
| Occupational exposure to animals (risk of zoonotic infections) | X | | | X | |
| History of IBS, IBD, chronic functional constipation, celiac disease, or other chronic gastrointestinal disorders | X | X | X | X | X |
| History of chronic systemic autoimmune diseases with gastrointestinal involvement | X | | X | X | X |
| History or high risk of gastrointestinal cancer or polyposis | X | X | X | X | X |
| Recent onset of diarrhea or hematochezia | X | | X | X | |
| History of neurological/neurodegenerative diseases | X | | X | X | |
| History of psychiatric disorders | X | | X | X | |
| Overweight/obesity | BMI ≥25 | | BMI >30 | BMI >25 | BMI >30 |
| Recent exposure to antibiotics, immunosuppressants, or chemotherapy | <3 months | Antibiotics | X | <3 months | <3 months |
| Chronic proton pump inhibitor use | X | | X | X | |
| Diarrhea (loose/liquid stools >3 times/day) | <6 months | | | X | <1 month |

| | | | | | |
|---|---|---|---|---|---|
| Recent illness or systemic symptoms (fever, sore throat, lymphadenopathy) | <2 weeks | Fever <2 weeks | | X | |
| Age limits | <18 or >50 | >60 | >60 | >60 | <16 or >60 |
| History of hepatitis A | | | X | | 6 months |
| Infection with multidrug-resistant bacteria | | | | | <3 months |
| Gastrointestinal surgery | X | | | | X |
| Metabolic syndrome, diabetes | X | X | | | X |
| Known history or risk factors for infectious diseases | | X | X | | |
| Recent hospitalization or stay in long-term care facilities | >2 days in last 3 months | | X | | |
| Recent needlestick injury | <6 months | X | ≤6 months | | |
| History of receiving growth hormone, bovine insulin, or coagulation factor concentrates | | Growth hormone | X | | |
| Known exposure or history of HIV-1/2, hepatitis B/C, syphilis, HTLV-1/2, malaria, Chagas disease, cutaneous tuberculosis, or mucosal herpes | X | X | | | |
| Risk factors for Creutzfeldt–Jakob disease (personal/family history, corneal transplant, cadaveric pituitary hormone use, bovine insulin use, nosocomial exposure, residence in UK/Ireland >3 months between 1980–1996 or >5 years in Europe since 1980) | X | X | | | |
| Sexual contact with individuals with HIV or viral hepatitis; men who have sex with men; partners of bisexual men | X | X | | | |
| New sexual partner within the last 12 months | X | X | | | |
| Moderate to severe malnutrition | X | | | | |

"X" indicates inclusion of the criterion; time restrictions are specified where applicable. Abbreviations: AST, aspartate aminotransferase; ALT, alanine aminotransferase; GGT, gamma-glutamyl transferase; CRP, C-reactive protein; HIV, human immunodeficiency virus; HTLV, human T-lymphotropic virus; CMV, cytomegalovirus; EBV, Epstein–Barr virus; PCR, polymerase chain reaction.
