## Supplementary Table 2 for "Identification of antibiotic resistance genes in fecal microbiota selected donors during the establishment of a biobank in the south of Brazil"

Supplementary Table 2. Laboratory screening workflow for donor eligibility

| Screening Type | Test Category | Specific Tests / Targets |
|---|---|---|
| **General Health (Blood Tests)** | Hematology | Complete blood count |
| | Electrolytes | Sodium, potassium, calcium, magnesium |
| | Renal function | Urea, creatinine |
| | Liver function | AST, ALT, GGT, total and fractionated bilirubin |
| | Metabolic/Nutritional markers | Glucose, vitamin B12, folic acid, vitamin D |
| | Inflammatory marker | C-reactive protein (CRP) |
| **Serological Screening (Blood-borne pathogens)** | Viral serology | HIV-1/2, Hepatitis A (IgM/IgG), Hepatitis B (HBsAg, anti-HBc IgG/IgM), Hepatitis C (anti-HCV), HTLV-1/2 |
| | Herpesviruses | Cytomegalovirus (CMV IgG/IgM), Epstein–Barr virus (EBV IgG/IgM), Herpes simplex virus I/II (IgG/IgM) |
| **Stool Microbiological Screening** | Conventional microbiology | Coproculture |
| | Toxin detection | *Clostridioides difficile* toxin A/B |
| **Molecular Screening (PCR-based assays)** | Targeted PCR | SARS-CoV-2, rotavirus, adenovirus |
| | Multiplex PCR panel (FilmArray®) | Bacteria: *Campylobacter spp.*, *Salmonella spp.*, *Shigella/EIEC*, *Vibrio cholerae*, *Yersinia enterocolitica*, *Clostridioides difficile*<br>Parasites: *Giardia lamblia*, *Entamoeba histolytica*, *Cryptosporidium*, *Cyclospora cayetanensis*<br>Viruses: Norovirus GI/GII, astrovirus, sapovirus, adenovirus F40/41 |

Abbreviations: AST, aspartate aminotransferase; ALT, alanine aminotransferase; GGT, gamma-glutamyl transferase; CRP, C-reactive protein; HIV, human immunodeficiency virus; HTLV, human T-lymphotropic virus; CMV, cytomegalovirus; EBV, Epstein–Barr virus; PCR, polymerase chain reaction.
